# FLARE: Fine-grained Learning for Alignment of spectra-molecule REpresentation Enhances Metabolite Annotation

**DOI:** 10.64898/2026.01.27.702086

**Authors:** Yan Zhou Chen, Blake Rushing, Soha Hassoun

**Author notes:** Contributing authors.

## Abstract

Accurate metabolite annotation via tandem mass spectrometry remains a major bottleneck in untargeted metabolomics. Recent implicit models that avoid molecular generation or spectra simulation have shown competitive performance by aligning spectra and molecular structures in the embedding space. Still, they overlook the detailed relationships between spectral peaks and molecular substructures that govern fragmentation. We introduce FLARE (Fine-grained Learning for Alignment of spectra-molecule REpresentations), a contrastive learning framework that leverages bidirectional peak-node alignment under learned weak supervision. Unlike models that rely solely on global embeddings, FLARE computes similarity via maxima over peak-to-atom and atom-to-peak interactions, capturing chemically meaningful local correspondences and enabling interpretable spectra–molecule matching. It achieves state-of-the-art results on MassSpecGym, with 43.15% rank@1 (mass-based) and 22.66% (formula-based), surpassing previous models by over 63%. FLARE’s learned embeddings correspond with molecular classes, match fingerprint similarity, and detect differential metabolites in a breast cancer xenograft study, showcasing its translational potential.

## 1 Introduction

Untargeted metabolomics plays a pivotal role in drug discovery, disease diagnosis, and the elucidation of biological mechanisms [1, 2]. These advances are largely enabled by tandem mass spectrometry (MS/MS), a technique that analyzes the chemical composition of complex samples. When coupled with liquid chromatography (LC), LC–MS/MS separates and fragments molecules, generating mass spectra that capture characteristic fragment ions. Each mass spectrum comprises the mass-to-charge ratios (m/z) of the charged fragments and their corresponding abundances. Molecular annotation is typically performed by matching experimental spectra to reference libraries. However, the coverage of these libraries remains limited due to the finite number of reference compounds and the sensitivity of fragmentation patterns to instrument settings, resulting in persistently low annotation rates [3]. These limitations reflect not only incomplete reference databases, but also the intrinsic difficulty of interpreting spectra generated by complex, stochastic physical processes.

A wide range of machine-learning (ML) methods have been developed in response to this challenge. These approaches can be broadly categorized as explicit or implicit. Explicit methods attempt to construct spectra from molecular structures or infer molecular structures directly from spectra [4–10] (Fig. 1a). Such methods have led to widely adopted tools, including SIRIUS [9] and CFM-ID [10]. However, by committing to predicting one modality from the other, explicit approaches necessarily discard complementary information contained in the unused modality and adopt a one-directional view of spectra–structure mapping relationships.

**Fig. 1:**
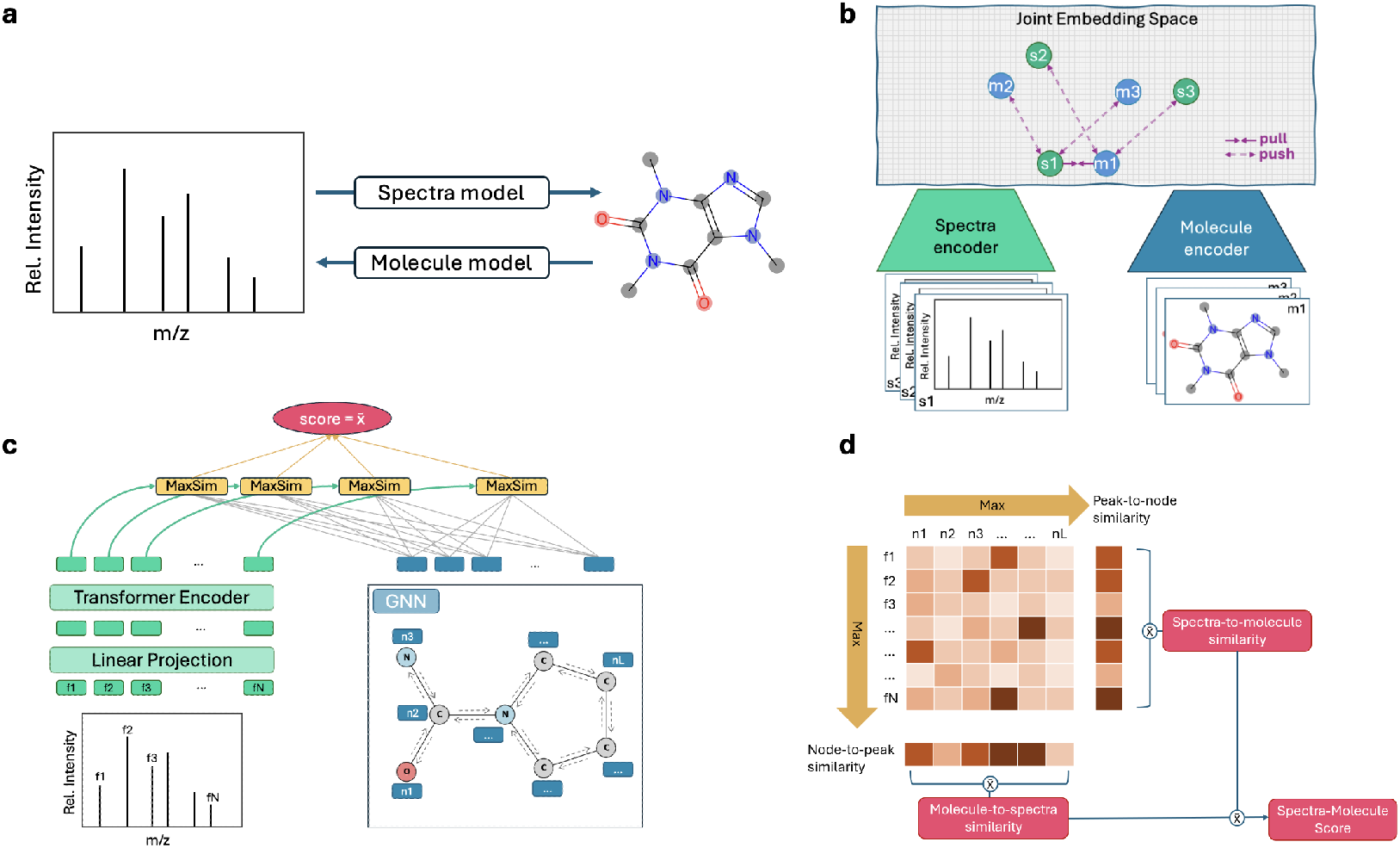
Annotation paradigms and overview of FLARE’s architecture and similarity computation methodology. **a**, Prior explicit methods construct one modality from the other, going from spectra to molecule or molecule to spectra. **b**, Implicit methods align spectra and molecules in a joint embedding space by effectively pulling matching pairs close together while pushing apart non-matching pairs. The push-pull interactions are only shown for one matching pair (s1, m1). **c**, FLARE encodes a spectra as a set of peak subformulae and a molecule as a graph. Peak representations are linearly projected and encoded with a transformer encoder. Node representations are embedded using a graph neural network (GNN). The spectra-molecule similarity is obtained by averaging the maximum node-to-peak similarities (not shown) and peak-to-node similarities. MaxSim is the maximum cosine similarity between two token embeddings. **d**, For a set of peaks f1,f2,…, fN and a set of nodes n1, n2, …,nL, peak-node similarities are computed using cosine similarity. Column-wise max yields node-to-peak similarity, and row-wise max yields peak-to-node similarity. To obtain molecule-to-spectra similarity, we average the node-to-peak similarities. Likewise, the average of peak-to-node similarities provides the spectra-to-molecule similarity score. The final spectra-molecule similarity score is computed as the average of molecule-to-spectra and spectra-to-molecule similarities.

Implicit approaches instead formulate spectra–structure annotation as a cross-modal alignment problem by embedding spectra and molecules within a shared representation space, where cosine similarities of the embeddings are used to rank candidate structures for a given spectrum [11–14] (Fig. 1b). These methods typically rely on contrastive learning frameworks inspired by CLIP [15] and its variants [16–18]. For example, JESTR [12] employs an InfoNCE objective [19] to maximize similarity between matched spectra–molecule pairs while separating non-matching pairs, and MVP [14] incorporates multiple spectral and molecular modalities to achieve state-of-the-art performance on the MassSpecGym benchmark [20]. Despite these advances, existing implicit methods primarily rely on global embedding alignment, even though fragmentation is governed by local, chemically constrained relationships between individual spectral peaks and molecular substructures. As a result, similarity scores computed from global representations are opaque, limiting the ability to validate learned structure or diagnose failure modes under distribution shift.

Implicit approaches expose a broader machine-intelligence problem: learning meaningful correspondences between heterogeneous modalities when the underlying relationships are latent, noisy, and never directly observed. In MS/MS, molecular structures encode latent physical and chemical constraints that govern stochastic fragmentation processes, giving rise to the observed spectral peaks. That is, the molecular graph encodes physical and chemical constraints, such as elemental composition and bond connectivity, that determine which fragmentation pathways are feasible, without explicitly specifying which peaks will be observed. The correspondence between peaks and molecular substructures is therefore not directly supervised and must therefore be inferred implicitly by any model that seeks to align spectra with molecular structures under substantial uncertainty arising from stochastic fragmentation and latent peak–substructure correspondence. In such settings, predictive performance alone is insufficient: without mechanisms to inspect and validate learned correspondences, it is difficult to assess whether a model has captured chemically meaningful structure or has instead exploited spurious correlations present in the data.

In parallel, the vision–language community has shown that fine-grained cross-modal alignment can improve both retrieval performance and interpretability. A prominent example is FILIP (Fine-grained Interactive Language–Image Pre-training), which extends CLIP by aligning individual word tokens with image patches and aggregating token–patch similarities via bidirectional maxima [16]. This formulation enables inspection of local correspondences that contribute to global similarity. However, vision–language alignment operates under semantic weak supervision, where tokens (words and image patches) are human-interpretable and semantically grounded. In contrast, spectra–structure alignment operates under physical weak supervision, where neither spectral peaks nor molecular fragments are directly observed, and incorrect alignments may violate chemical feasibility rather than semantic coherence. This raises a fundamental machine-intelligence question: can fine-grained contrastive alignment operate effectively under physical weak supervision and provide evidence about what a model has learned when local correspondences are latent and uncertain?

Building on these insights, we introduce FLARE (Fine-grained Learning for Alignment of spectra–molecule REpresentations), a contrastive learning framework designed to learn fine-grained spectra–structure correspondences under physical weak supervision. FLARE replaces global embedding similarity with a learning objective based on bidirectional peak–atom alignment, encouraging representations in which spectra–structure compatibility emerges from local, mutually consistent correspondences. By leveraging graph neural networks (GNNs) [21] to learn context-aware atom-level representations, FLARE grounds peak–atom alignment in chemically meaningful sub-structural constraints. Crucially, this formulation enables alignment hypotheses to be inspected and evaluated against chemical plausibility, providing a mechanism to validate learned representations under uncertainty, rather than treating similarity scores as uninterpretable latent metrics.

FLARE’s learning formulation translates into strong empirical evidence across both benchmarked and real-world settings. On the MassSpecGym retrieval benchmark, FLARE achieves state-of-the-art performance when retrieving candidate structures by mass and by formula, demonstrating that fine-grained alignment does not compromise global discriminative capability. Beyond retrieval accuracy, the learned joint embedding space organizes spectra according to molecular classes and structural similarity, indicating that the model internalizes chemically meaningful organization rather than relying on spurious global correlations. In contrast to global alignment approaches such as JESTR, which optimize a single similarity score between pooled representations, FLARE induces a complementary learning bias by shaping representations through local, bidirectional peak–substructure correspondences that can be directly inspected and evaluated. Finally, in a case study using patient-derived xenograft tumors from metastatic and non-metastatic breast cancers, FLARE identifies differential spectral features that distinguish tumor phenotypes, illustrating how fine-grained alignment supports validation and biological insight in realistic discovery settings.

FLARE is released as an open-source framework and includes an interactive visualization tool for exploring learned peak–substructure correspondences, publicly available on Hugging Face (https://huggingface.co/spaces/HassounLab/FLARE).

## 2 Results

### 2.1 FLARE overview

FLARE aligns tandem mass spectra with molecular structures under physical weak supervision. The method is built around the idea that spectra–structure compatibility is best understood as the accumulation of local, mutually consistent evidence between spectral peaks and molecular substructures, rather than as a property of pooled global representations.

Rather than collapsing spectra or molecules into single vectors, FLARE represents each modality as a collection of local element: spectral peaks and molecular nodes that can participate directly in alignment. To ensure that these local elements are informative, FLARE encodes contextual structure within each modality prior to alignment. Atom-level representations are learned using graph neural networks, allowing each atom to aggregate information from its bonded neighbors and reflect chemically meaningful substructures. Spectral peaks are similarly embedded with contextual information that captures their spectral characteristics. Alignment therefore, operates over context-aware embeddings that represent structured molecular regions and coherent spectral features, rather than over embeddings of isolated peak or atom tokens.

Spectra–structure compatibility in FLARE is computed through a bidirectional alignment mechanism (Fig. 1c, and 1d). In one direction, individual spectral peaks are matched to regions of the molecular graph; in the other, molecular substructures are evaluated based on the spectral evidence supporting them. Treating these two directions symmetrically enforces a mutual-consistency constraint, ensuring that compatibility is supported from both spectral and structural perspectives. Local alignment scores are then aggregated to produce a global similarity score for a spectrum–molecule pair. Importantly, this score is derived from the collection of local correspondences, rather than computed directly from pooled embeddings. Global compatibility therefore reflects the cumulative support of fine-grained evidence across the two modalities. FLARE is trained using paired spectra–molecule examples without any supervision at the level of peak–substructure correspondences. The learning objective encourages correct global pairing by shaping local alignments, forcing the model to infer latent correspondences consistent with both modalities.

A key consequence of FLARE’s learning formulation is inspectability. Because global similarity is derived from collections of local alignments, the contributions of individual peaks and molecular substructures to spectra–structure compatibility can be examined directly. This provides a mechanism to assess whether learned representations reflect chemically plausible relationships, offering a means to validate learning behavior under substantial uncertainty. In contrast to global contrastive approaches, where similarity scores are computed from pooled embeddings with opaque internal structure, FLARE exposes the intermediate correspondences that drive alignment. The FLARE architecture is described in detail in the Methods section of the paper.

### 2.2 Fine-grained alignment achieves state-of-the-art retrieval performance

We first evaluate whether fine-grained spectra–structure alignment can support accurate molecular retrieval. We assess FLARE on the MassSpecGym benchmark [20], a standardized evaluation suite for spectra–molecule retrieval that reflects realistic levels of chemical diversity and annotation uncertainty (Sec. 4.1). Retrieval is performed under commonly used candidate selection settings, including retrieval by precursor mass and by molecular formula. Across all evaluation metrics (Table 1), FLARE outperforms previous approaches when retrieving candidates by mass and formula. FLARE achieves the highest rank@1 accuracy and related evaluation measures among all compared methods, establishing state-of-the-art performance on MassSpecGym. Examining the results, we note that contrastive-based models (JESTR, MVP, and FLARE) that leverage cross-modal interactions achieve notably stronger performance than one-directional models (MIST and ESP). Compared to MIST, which also represents peaks as subformulas, FLARE improves rank@1 by 194.74% and MCES@1 by 48.99% for mass-based retrieval and 136.77% and 29.41%, respectively, for formula-based retrieval. Relative to previous contrastive-learning models, FLARE achieves further substantial gains. It surpasses the second-best model, MVP, which benefits from additional spectral and molecular views during training, by 63.63% in rank@1 and 36.10% in MCES@1 for mass-based retrieval and 104.14% and 22.48%, respectively, for formula-based retrieval. These substantial improvements demonstrate that replacing global embedding similarity with fine-grained, bidirectional alignment not only preserves discriminative capability but also leads to improved retrieval accuracy.

**Table 1:** Ranking performance on MassSpecGym compared to other tools. The best performance is highlighted, and the second best performance is underlined.

| Model | rank@1 (↑) | rank@5 (↑) | rank@20 (↑) | MCES@1(↓) |
| --- | --- | --- | --- | --- |
| A. Candidates by Mass |  |  |  |  |
| MIST | 14.64 | 34.87 | 59.15 | 15.37 |
| ESP | 10.71 | 24.84 | 42.66 | 35.36 |
| JESTR | 15.13 | 36.75 | 60.32 | 20.71 |
| MVP | <u>26.37</u> | <u>58.90</u> | <u>86.88</u> | <u>12.27</u> |
| FLARE | **43.15** | **75.59** | **92.89** | **7.84** |
| B. Candidates by Formula |  |  |  |  |
| MIST | 9.57 | 22.11 | 41.12 | 12.75 |
| ESP | 11.05 | 27.42 | 52.20 | 16.44 |
| JESTR | <u>11.85</u> | <u>32.95</u> | 61.46 | 11.68 |
| MVP | 11.10 | 31.14 | <u>61.99</u> | <u>11.61</u> |
| FLARE | **22.66** | **50.00** | **75.15** | **9.00** |

### 2.3 Fine-grained alignment induces a structured representation space

The fine-grained alignment of spectra and molecules in FLARE encourages the emergence of a unified embedding space across modalities. A chemically meaningful embedding space should reflect the underlying chemical structure in a systematic and interpretable manner. Here, we show that fine-grained alignment learned under weak supervision induces a chemically organized embedding space that exhibits monotonic correspondence with chemical similarity, structured uncertainty, cross-modal consistency, and semantic coherence.

We first examine whether the organization of the learned spectral embeddings manifests as a semantically coherent structure by visualizing the spectral embedding space (Sec. 4.5). UMAP visualization of spectral embeddings colored by chemical class reveals contiguous regions corresponding to major chemical classes (Fig. 2a), despite the absence of class-level supervision. Some subclasses, such as alkaloids, appear intermingled in two dimensions, potentially reflecting chemical diversity and projection limitations rather than the absence of structure.

**Fig. 2:**
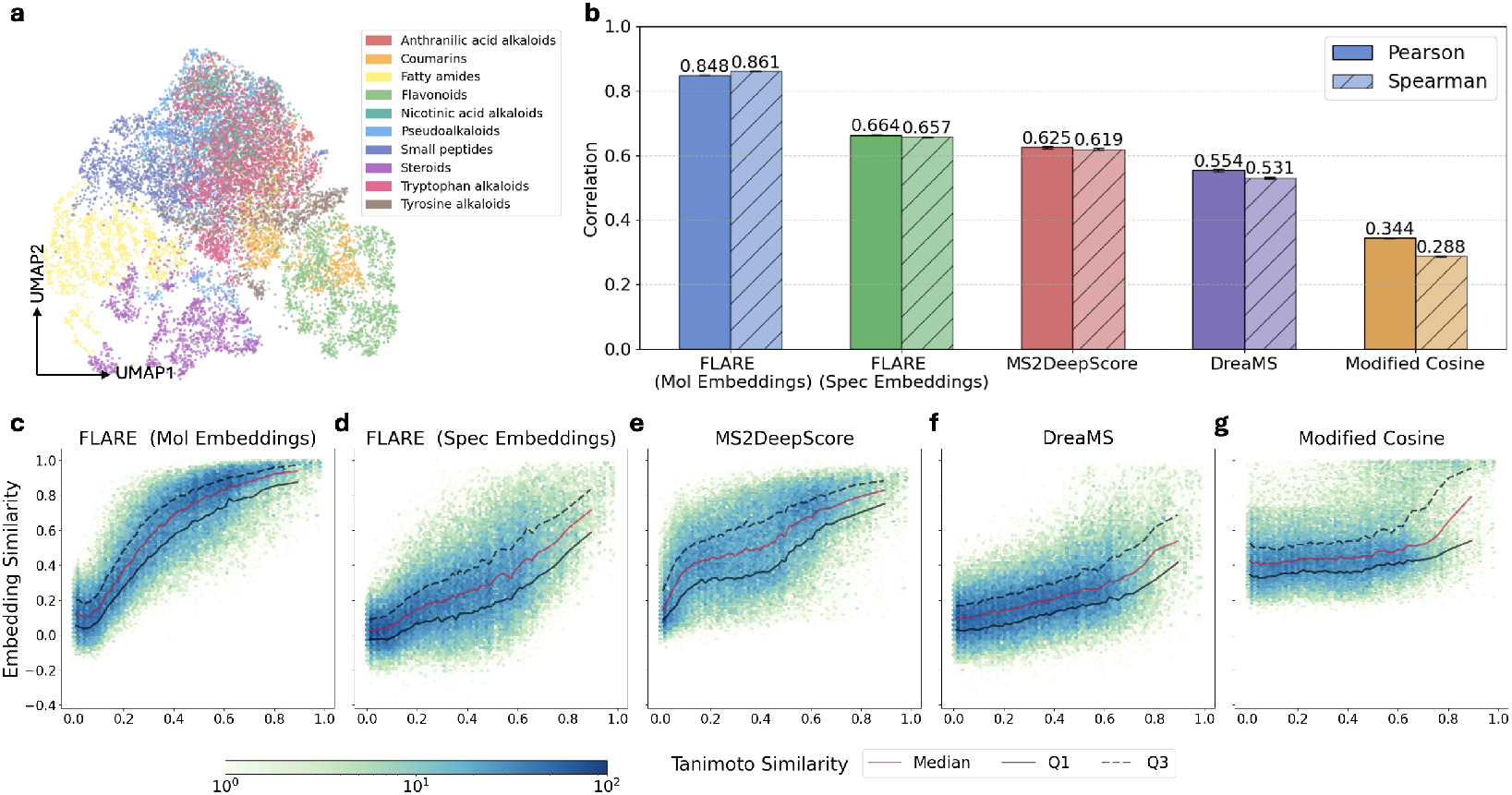
Evaluation of the learned embedding space. **a**, UMAP projection of MSGym spectral embeddings colored by chemical superclasses demonstrates biologically meaningful clustering. Only a subset of spectra from the ten most frequent classes is shown. **b**, Average Pearson and Spearman correlations between embedding similarity and chemical similarity (Tanimoto) across 5 independently repeated runs (standard deviations are shown in the error bar and appear as short dashed lines at the top of each bar). **c-g**, Hexbin plot of embedding similarity against fingerprint Tanimoto similarity using FLARE molecule embeddings, FLARE spectral embeddings, MS2DeepScore, DreaMS, and modified cosine similarity.

Next, we evaluate the relationship between embedding similarity and fingerprint similarity using over 50,000 molecule pairs. The experiment is repeated 5 times, and consistent trends are observed (Sec. 4.4). We first assess whether embedding similarity increases monotonically with chemical similarity. To quantify this relationship, we compute the average Pearson and Spearman correlation coefficients across all 5 experiments (Fig. 2b). FLARE’s molecular embeddings show highest correlations (Pearson 0.848; Spearman 0.861), as expected given their direct access to molecular structure. The spectral embeddings also exhibit a strong monotonic relationship (Pearson 0.664; Spearman 0.657), indicating that fine-grained alignment yields representations in both modalities that independently preserve molecular similarity structure. Compared to MS2DeepScore [22, 23], which is trained explicitly to predict Tanimoto similarity from spectral pairs, FLARE exhibits stronger monotonic association, while DreaMS [24], a large unsupervised spectral model, shows weaker correspondence. FLARE, among other embedding-based method, show substantially stronger correspondence compared to modified cosine similarity, a standard similarity measure for matching reference spectra [25].

Monotonic correspondence captures average behavior but obscures variability across individual molecule pairs. To characterize this heterogeneity, we analyze the dispersion of embedding similarity conditioned on chemical similarity using the first and third quartiles (Q1 and Q3) relative to the median for one of the experiments (Fig. 2c-g). FLARE’s molecular embeddings display markedly low dispersion, evidenced by the tight Q1 and Q3 bounds, as they bypass fragmentation variability by encoding molecular structure directly (Fig. 2c). Spectral-only FLARE exhibits increased dispersion at high Tanimoto similarity, reflecting intrinsic fragmentation variability, mismatches between fingerprints and fragmentation-relevant features, and the inductive bias of fine-grained alignment, which aggregates heterogeneous local correspondences rather than enforcing a deterministic similarity mapping (Fig. 2d). MS2DeepScore displays reduced dispersion at high Tanimoto similarity, consistent with objective-driven compression toward the training target, whereas DreaMS exhibits broader variability across regimes, resembling the behavior of FLARE’s spectral embeddings (Fig. 2e and 2f). Importantly, the increased dispersion observed for FLARE does not indicate a loss of global organization: spectral embeddings preserve strong rank ordering while faithfully representing ambiguity inherent to spectra–structure relationships, as is the case with modified cosine similarity (Fig. 2g).

Finally, we visit whether spectral and molecular representations organize chemical space consistently. Cross-modal consistency here refers to whether both modalities independently reflect the same underlying notion of chemical similarity, rather than direct similarity between embeddings. Consistent with the Pearson and Spearman correlations, Figures 2c and 2d illustrate monotonic trends across similarity regimes for both spectral and molecular embeddings, confirming that the observed correlations are not localized to a particular Tanimoto similarity range. Importantly, the shared organization emerges without explicit supervision, enforcing agreement between modalities, indicating that fine-grained alignment induces a common chemical organizing principle while preserving modality-specific information. Together, these observations show that fine-grained alignment yields spectral representations that are chemically grounded, uncertainty-aware, and semantically coherent.

### 2.4 Fine-grained contrastive vs conventional global-based contrastive

Fine-grained and global contrastive learning differ fundamentally in how they define spectra–structure compatibility. To examine how these differences manifest in practice, we conducted a systematic comparison between FLARE, which employs fine-grained peak–node alignment, and JESTR, which uses conventional global contrastive learning. Both models are evaluated on the MassSpecGym test set with candidates by formula.

We first examine how the two models differ in their ability to prioritize the correct molecular structure by analyzing the ranking position assigned to the target molecule for each test spectrum (Fig. 3a). This comparison shows that FLARE more frequently ranks the correct (target) molecule higher than JESTR (FLARE wins). At the same time, there exists a non-trivial subset of spectra where JESTR substantially outperforms FLARE (JESTR wins), suggesting that the two models capture partially distinct aspects of molecular-spectra compatibility.

**Fig. 3:**
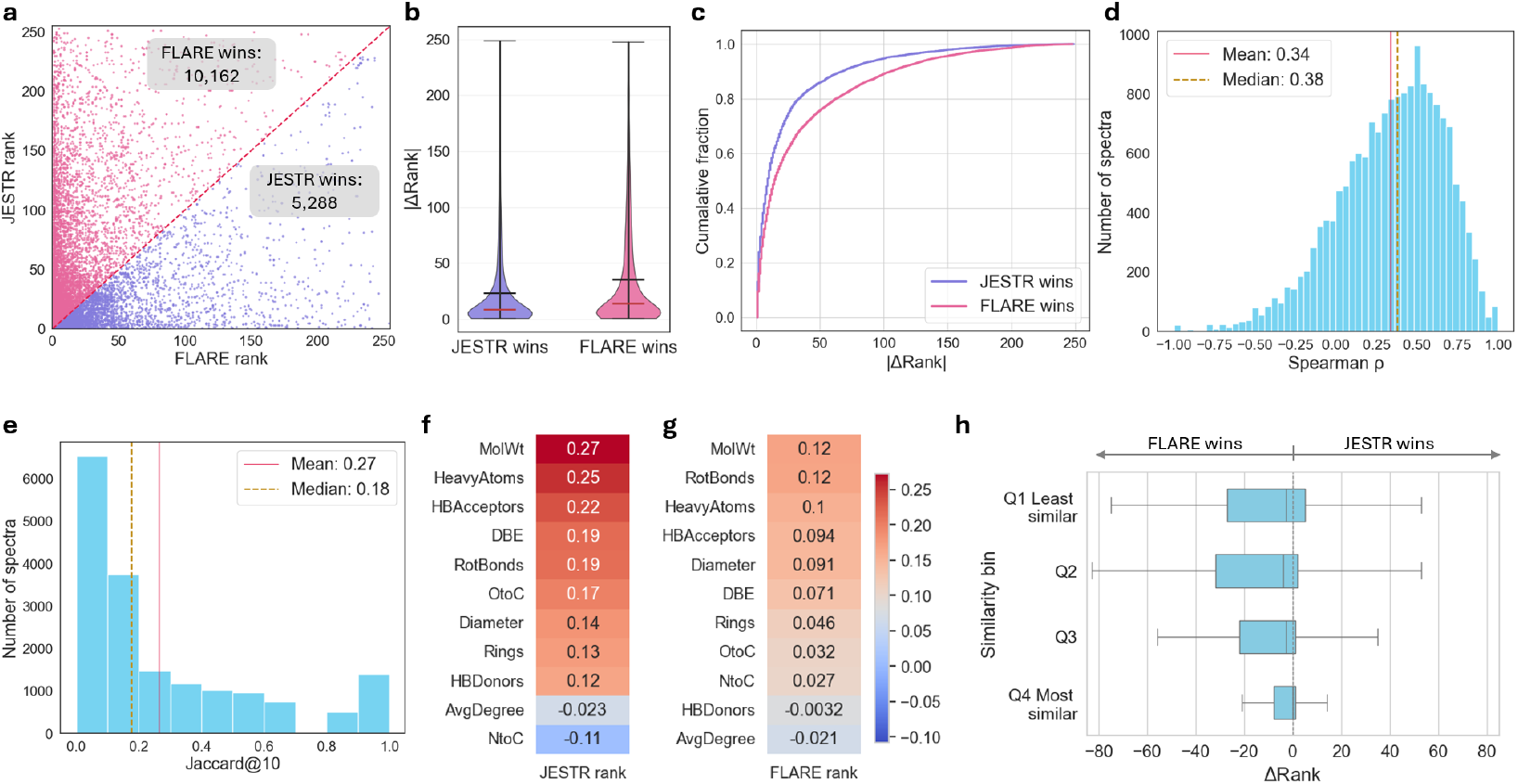
Comparison between fine-grained contrastive and conventional global-based contrastive ranking results. **a**, Comparison of the target molecule rank computed by JESTR and FLARE. **b**, Distribution of the rank difference for instances where JESTR wins (i.e., JESTR rank *<* FLARE rank), and where FLARE wins (i.e., FLARE rank *<* JESTR rank). **c**, Cumulative fraction of JESTR wins vs FLARE wins by rank difference. **d**, Distribution of Spearman *ρ* coefficients for each spectrum and its candidate rank orders by JESTR and FLARE. **e**, Distribution of the Jaccard index for the top-10 candidates from JESTR and FLARE. (**f, g**), Inverse correlation between molecular descriptors and JESTR or FLARE rank. **h**, boxplots of rank differences categorized by similarity bins. Similarity is determined by the maximum similarity between the target molecule and training molecules.

To quantify the magnitude of these differences, we analyze the distribution of absolute rank differences ( |Δ Rank |) for spectra where each model wins. FLARE’s wins tend to occur with larger rank improvements, as reflected by the wider upper tail in the violin plot and the slower rise of the cumulative fraction curve, whereas JESTR’s wins are more concentrated at smaller rank differences (Fig. 3b and 3c). Together, these results demonstrate that the fine-grained similarity score provides a pronounced advantage in discriminating the correct structure from a pool of candidates.

Next, we assess the degree of agreement for a given spectrum and its candidates. For each spectrum, we compute the Spearman correlation of candidate ranks output by FLARE and JESTR. The distribution of the Spearman coefficients shows a weak positive (mean: 0.34; median: 0.38) correlation, reflecting partial agreement, but considerable differences (Fig. 3d). Within the top 10 candidates, most spectra have a low Jaccard index (mean: 0.27; median 0.18) between the two models, which further confirms that each model emphasizes distinct spectral-molecular comparability features (Fig. 3e).

To better understand what underlies the observed rank differences, we analyze whether model performance can be explained by simple, interpretable molecular descriptors. Specifically, we evaluate a set of molecular descriptors (Table S3) and measure their inverse correlation with target numerical rank. Across all descriptors, no strong correlations are observed for either model, suggesting the performance difference is not well explained by coarse molecular properties alone (Fig. 3f and 3g). The strongest trend appears in JESTR, which exhibits a weak positive correlation (*r* = 0.27) between molecular weight (MolWt) and rank performance, indicating relatively better performance on larger molecules. FLARE, in contrast, shows a weaker correlation with molecular weight (*r* = 0.12) and consistently low correlations across all other descriptors. The absence of strong descriptor-level correlations for FLARE suggests that its performance gains arise from more localized chemically specific interactions captured through peak-node level embedding alignment, rather than from sensitivity to global molecular characteristics. Consequently, we expect FLARE to exhibit stronger generalization, particularly when test molecules differ substantially from those seen during training.

To test this hypothesis, we group test molecules by their maximum Tanimoto similarity to training molecules and analyzed the corresponding Δrank distributions (Fig 3h). Molecules are split by quartiles, where Q1 corresponds to the 25th percentile of similarity (least similar to training molecules) and Q4 to the 75th percentile (most similar to training molecules). For highly similar molecules, performance differences between FLARE and FLARE are relatively small, with narrow Δrank distributions centered close to zero, indicating comparable ranking behavior. For less similar molecules, FLARE increasingly outperforms JESTR by larger margins, as evidenced by the leftward shift of the Δrank distributions. This trend confirms that fine-grained alignment is particularly beneficial when generalizing beyond the training distribution, enabling more robust alignment of novel spectra and molecules.

Overall, these analyses demonstrate that fine-grained peak–node alignment provides a substantial advantage in prioritizing the correct molecular structure, particularly in challenging settings where candidate molecules are structurally diverse or dissimilar to the training distribution. FLARE not only wins more often, but does so by larger margins, reflecting its ability to leverage localized, chemically specific correspondences that are not captured by global contrastive objectives. At the same time, the consistent presence of cases in which JESTR outperforms FLARE underscores the value of global similarity, which can be advantageous for certain spectra, especially for larger molecules and those that closely resemble training molecules. The weak agreement between the two models further highlights that they encode complementary notions of spectra–structure compatibility. Importantly, no simple molecular descriptors or similarity thresholds reliably predict when one paradigm will dominate, suggesting that the strengths of fine-grained and global similarity are context dependent. These findings motivate future hybrid models that explicitly integrate and prioritize both fine-grained and global similarity signals, which may yield further improvements.

### 2.5 Inspectable peak–substructure correspondences validate learning under uncertainty

FLARE’s fine-grained alignment architecture produces similarity scores that are explicitly decomposable into local peak–substructure interactions, providing a direct inspection of how spectral evidence supports molecular compatibility. Rather than deriving similarity from pooled global embeddings, FLARE computes spectrum–molecule compatibility through bidirectional interactions between spectral peak embeddings and molecular node embeddings. Visualization of the resulting peak–node similarity matrix reveals individual correspondences between spectral peaks and molecular substructures, offering an interpretable account of the model’s decision-making process.

We analyze representative cases to assess whether these learned correspondences are chemically meaningful and to characterize failure modes. To evaluate chemical plausibility, we annotate each peak with the most likely fragment suggested by MAGMa,[26], which iteratively fragments candidate molecules to explain observed spectral peaks. When FLARE correctly ranks the target molecule highly, peak–substructure alignments generally conform to established fragmentation behavior. For example, a peak at m/z 128.0277, annotated as C4H5N3S, aligns most strongly with the corresponding fragment assigned by MAGMa. Similarly, the peak at m/z 167.9939 (C7H5NS2) exhibits a strong correspondence with a chemically compatible substructure in the molecular graph (Fig. 4a). In these cases, high global similarity emerges from the aggregation of multiple localized, chemically plausible correspondences rather than reliance on a single dominant feature.

**Fig. 4:**
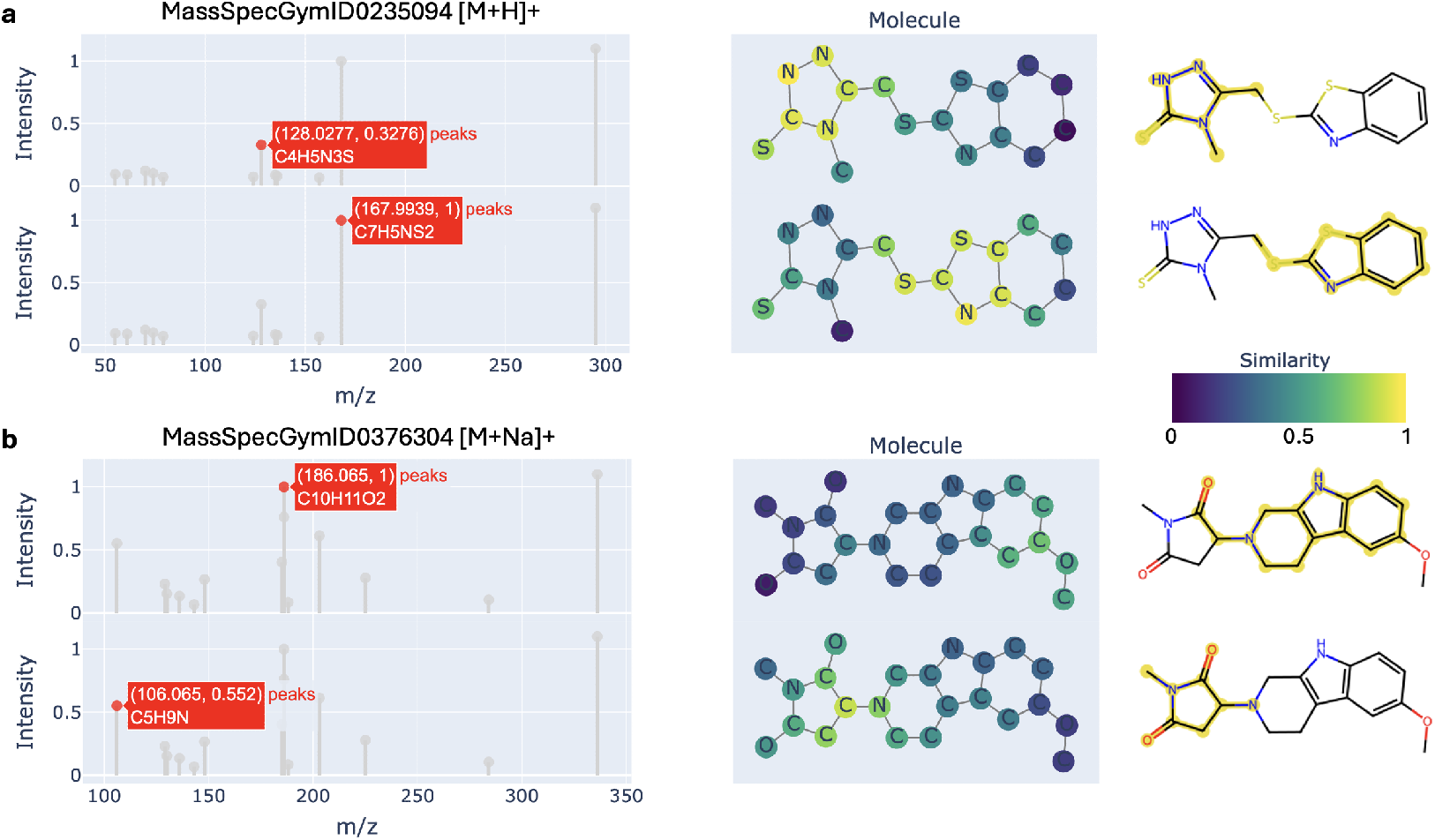
Peak–substructure alignment visualization. From left to right: a selected spectral peak, molecular graph nodes colored by similarity to the corresponding peak embedding, and the most likely fragment proposed by MAGMa. **a**, MassSpecGymID0235094 illustrates a successful case in which the target molecule is ranked correctly, driven by strong and chemically plausible correspondences between spectral peaks and molecular substructures. **b**, MassSpecGymID0376304 shows a failure case in which the target molecule is incorrectly ranked, accompanied by diffuse and chemically inconsistent peak–substructure alignments relative to plausible fragmentation patterns.

By contrast, spectra for which FLARE performs poorly display chemically inconsistent alignment patterns, often arising from incorrect or ambiguous subformula annotations. For instance, the peak at m/z 186.065 is assigned the formula C10H11O2. However, generating such a fragment would require extensive rearrangements of the precursor molecule while simultaneously eliminating all nitrogen atoms, a process inconsistent with common fragmentation mechanisms. This uncertainty is reflected in the model’s diffuse similarity across molecular nodes (Fig. 4b). In this case, MAGMa proposes a fragment with the formula C13H11N2O instead. In another example, a peak at m/z 106.065 (C5H9N) aligns with a plausible substructure, yet would imply the loss of two oxygen atoms, contradicting the fragment suggested by MAGMa.

These discrepancies arise in part from the subformula procedure, which uses a naive combinatorial strategy that selects the formula whose mass most closely matches the observed m/z while assuming the presence of the adduct in each fragment. While more sophisticated formula assignment methods could improve alignment quality, the current strategy is computationally efficient and scalable. Moreover, exposure to a large number of examples enables the model to learn robust alignment patterns despite this source of noise.

Taken together, these analyses demonstrate that FLARE’s fine-grained alignment provides more than improved retrieval accuracy. By exposing the internal evidence underlying spectra–structure compatibility, FLARE enables users to assess whether predictions are supported by chemically plausible peak–substructure correspondences. This inspectability facilitates the validation of learned representations under weak supervision and structured uncertainty, distinguishing FLARE from global similarity models whose predictions cannot be decomposed or interrogated. In practice, such interpretability offers a powerful diagnostic tool for understanding model behavior, identifying annotation errors, and building trust in spectral annotation workflows.

### 2.6 FLARE identifies differential metabolites in metastatic and non-metastatic breast cancer

To evaluate FLARE in a realistic biological setting, we applied FLARE to patient-derived xenograft (PDX) models of breast cancer to identify metabolomic differences that distinguish metastatic from non-metastatic (Primary) tumors. Beyond assessing annotation accuracy, this case study probes whether FLARE can support meaningful biological analysis in regimes characterized by incomplete annotations, heterogeneous spectra, and substantial uncertainty—conditions that are typical of real-world metabolomics experiments. Sample preparation and feature extraction are described in Sec. 4.8.

The dataset comprises nine metastatic and eight primary tumors, from which 18,772 features were extracted, including 13,594 features associated with MS/MS spectra. Using SIRIUS v6.1.0, 5,799 spectra were assigned precursor formulas and adducts; most of the remaining spectra carried unsupported charge states (*>*+1). Candidate structures were retrieved by mass (0.01 Da tolerance) from HMDB[27], COCONUT[28], a curated database of biologically relevant molecules[29], as well as molecular entries derived from multiple spectral libraries, including MassSpecGym, NIST23[30], and the GNPS MULTIPLEX dataset[31].

We first evaluate FLARE’s performance on annotating observed spectra based on a relevant in-house reference library of standards. Among 5,562 spectra with candidates, 116 had a match to an in-house standard based on the spectral similarity, neutral mass error, and retention time error. The correct molecular structure appeared among the candidates for 74 spectra, and FLARE ranked the true structure first for 29 spectra, yielding a rank@1 accuracy of 39.19%, consistent with the reported performance on the MassSpecGym benchmark. When restricting analysis to spectra for which SIRIUS assigned the correct precursor formula (66 cases), FLARE achieved a rank@1 accuracy of 43.9% (29/66), indicating that performance in this real-world setting aligns with controlled benchmark results.

We next asked whether FLARE-derived annotations support biologically meaningful differential analysis. Using high-confidence annotations (matching score *>* 0.35), we performed differential abundance analysis between metastatic and primary tumors. A volcano plot identified 54 differential features (p ≤ 0.05 and |log2 fold change| ≥ 1) (Fig. 5a). A clustered heatmap of the 50 most significant features identified two major sample groups (Fig. 5b). Notably, four metastatic tumor samples clustered with all primary tumors, suggesting heterogeneity within the metastatic group and potentially reflecting different stages of metastatic progression.

**Fig. 5:**
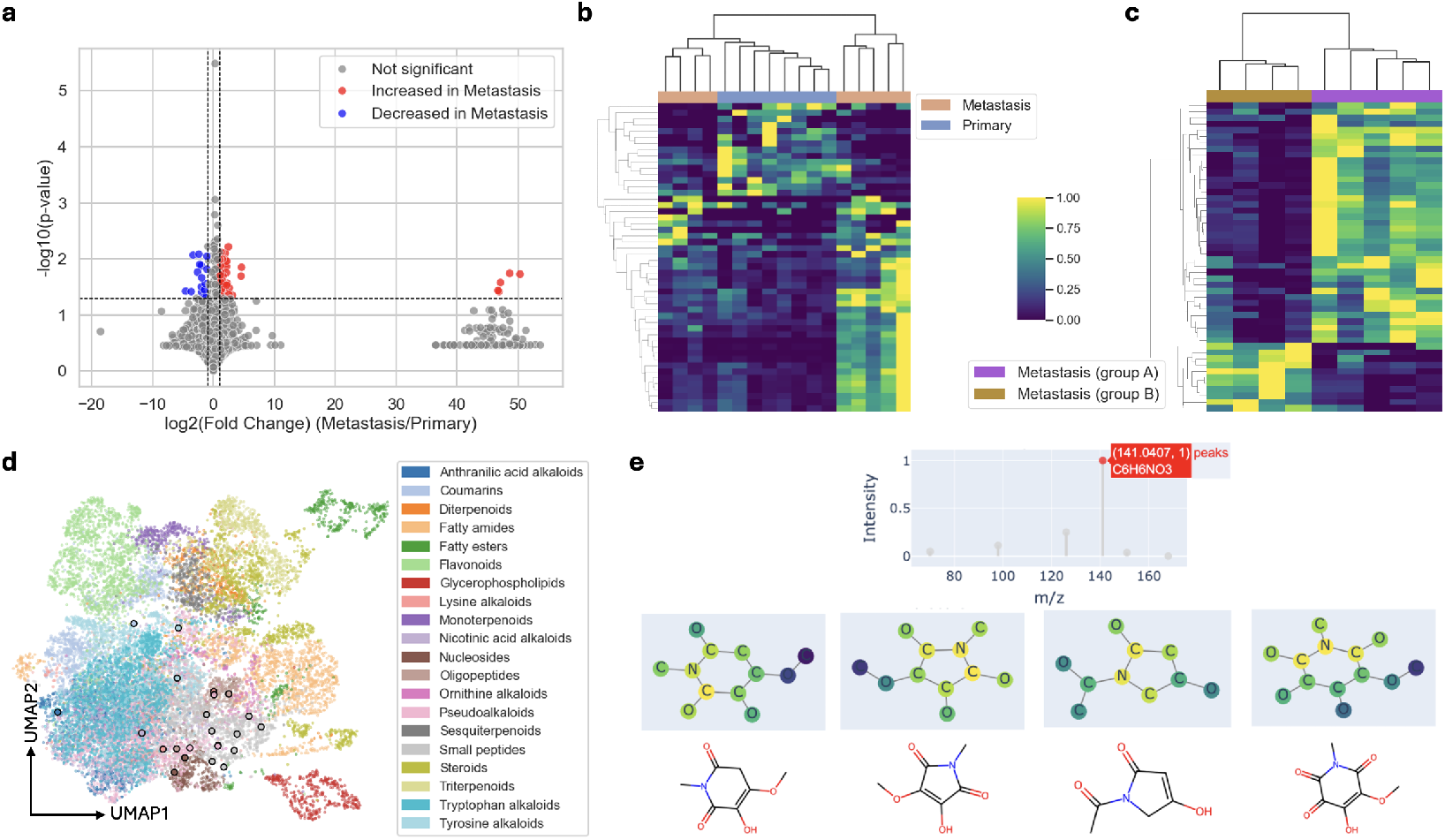
Metastatic breast cancer dataset analyses. **a**, A volcano plot of feature abundance comparing metastasis samples and primary samples, identifying 54 differential features. **b**, A clustered heatmap using the 50 most significant differential features results in two major clusters. **c**, A secondary differential analysis comparing the two groups of metastasis identifies 88 significant features, and the 50 most significant features result in two distinct clusters when used in a clustered heatmap analysis. **d**, UMAP projection of a subset of MassSpecGym spectra colored by superclasses and 23 significant features with low confidence annotation colored by the predicted superclass and outlined in black. **e**, Peak-to-node visualization of a spectrum corresponding to a significant feature with low confidence annotation. Candidates are identified from a set of 2 million molecules based on the embedding similarity. The correspondences suggest possible substructures within the target molecule.

To further interrogate the two separate groups of metastasis, we repeated the differential analysis restricted to metastatic tumors, identifying 88 differential features. The corresponding clustered heatmap revealed pronounced metabolic differences between the two metastatic subtypes (Fig. 5c). When reclustering all samples using the same feature set, three primary tumors grouped with the Metastasis group B samples, while five primary tumors remained in a distinct cluster (Fig. S1). These patterns suggest that a subset of primary tumors shares metabolic features with metastatic tumors, potentially marking early or transitional states of metastatic progression.

A major challenge in untargeted metabolomics is the large fraction of spectra that lack confident molecular annotations. In this dataset, 1,653 spectra did not meet the high-confidence annotation threshold. Rather than discarding these features, we examined whether FLARE could still extract biologically informative structure from this subset. A volcano plot comparing metastatic and primary tumors identified 23 differential features within this subset. Using k-nearest neighbors, we predicted the classes of these 23 spectra and embedded them into the UMAP space alongside a subset of MassSpecGym reference spectra (Sec. 4.6, Fig. 5d). We then apply both the spectral and molecular encoders to retrieve the five most similar molecular candidates for each feature within the embedding space, hence providing candidate-level hypotheses despite the absence of definitive annotations (Sec. 4.7).

Crucially, FLARE enables inspection of peak–substructure correspondences for these partially characterized features. For example, a spectra contains a peak at m/z 141.0407 that shows strong similarity to nodes corresponding to all or part of the subformula C6H6NO3 in several candidate molecules (Fig. 5e). These localized correspondences provide interpretable evidence for plausible substructural elements even when full molecular identification is not possible.

This case study demonstrates that FLARE supports metabolomic analysis beyond conventional annotation-first pipelines. By combining chemically organized embeddings with inspectable peak–substructure correspondences, FLARE enables differential analysis, subgroup discovery, and interpretable hypothesis generation under conditions of weak supervision and structured uncertainty. These capabilities are essential for real-world metabolomics, where incomplete annotations and ambiguous spectra are the norm rather than the exception.

## 3 Conclusion

We present FLARE, the first implicit metabolite annotation framework to incorporate fine-grained spectra–structure alignment, enabling peak-to-substructure interpretability within a contrastive learning paradigm. By bidirectionally maximizing similarity between peak- and atom-level embeddings, FLARE captures chemically meaningful peak-to-fragment relationships that conventional global methods overlook. This design yields substantial performance gains on the MassSpecGym benchmark and produces a chemically organized joint embedding space suitable for downstream similarity prediction and molecular class inference.

Beyond retrieval accuracy, FLARE enables transparent inspection of spectra–structure compatibility through peak-to-node similarity maps, allowing practitioners to diagnose fragmentation ambiguities, identify unreliable subformula assignments, and gain mechanistic insight into model predictions. Our systematic comparison with a global contrastive model reveals that fine-grained and global strategies capture complementary aspects of spectral and structural information, suggesting promising directions for hybrid architectures that unify local and global cross-modal signals.

In a case study of metastatic breast cancer, FLARE identifies differential metabolic features and provides partial structural annotation when alignment scores are low, demonstrating its utility in real-world discovery settings where full structural identification is often infeasible. Overall, FLARE establishes an interpretable, high-accuracy foundation for metabolite annotation, setting a new standard for reliable machine-learning approaches in metabolomics and chemical discovery.

## 4 Methods

### 4.1 Dataset

We evaluate FLARE on the retrieval task using the MassSpecGym benchmark (Fig. S1). To evaluate retrieval, each spectrum in this dataset is associated with two molecular candidate sets, retrieved either by mass (within a 10 ppm tolerance) or by molecular formula. MassSpecGym provides a challenging test split in which all test molecules are separated by at least 10 MCES[32] distance from every molecule in the training and validation sets. Strong performance on this benchmark demonstrates the model’s generalizability.

### 4.2 Input Representation and Encoders

FLARE takes a spectrum and a molecule as input, assuming that the parent chemical formula and adduct are known. We adopt the spectrum and molecule featurizers used in MVP. For spectral processing, peaks are first assigned subformulae with a tolerance of *±*20 ppm using the enumeration-based algorithm from MIST. To reduce noise, only the 60 most intense peaks are retained. Each subformula is encoded as a 14-dimensional vector, where each dimension corresponds to the count of an element *e* ∈ *E, E* = {*C, H, O, N, P, S, Cl, F, Br, I, B, As, Si, Se*}. The subformula encoding is normalized by dividing each element count by the maximum observed in the training dataset. The resulting subformula encoding is concatenated with the normalized peak intensity to yield the final peak-level representation. The parent formula is explicitly included in all spectra and assigned a fixed normalized intensity of 1.1, which empirically improved model performance.

Each peak is first projected through a shallow multilayer perceptron (MLP) to obtain a dense latent representation. These embeddings are subsequently processed by a transformer encoder, which models contextual relationships among peaks and produces a set of peak embeddings.

Molecular structures are represented as undirected graphs, with atoms as nodes and bonds as edges. Node feature vectors comprise atom type, atomic mass, valence, ring membership, formal charge, number of radical electrons, chirality, degree, hydrogen count, and aromaticity indicators. The molecular encoder is implemented as a standard graph convolutional network (GCN)[33], which iteratively propagates and aggregates local neighborhood information to derive node-level embeddings.

Detailed architectural specifications, parameter values, and hyperparameter search ranges are provided in Table S2.

### 4.3 Fine-grained Contrastive Learning

We adopt a standard contrastive learning framework, but modify the similarity (discriminator) function using fine-grained matching inspired by FILIP. Instead of aggregating spectra peaks and molecular nodes into global embeddings, we compute similarity between spectra peaks and molecule nodes.

Given a spectrum *s* with *N* peaks and a molecule *m* with *K* nodes, we embed the spectra as *f*_*θ*_(*s*) ∈ ℝ^*N×d*^, and the molecule as *g*_*ϕ*_(*m*) ∈ ℝ^*K×d*^. For spectra-to-molecule similarity, each peak is compared to all nodes by computing the cosine similarity and retaining the maximum. Finally, we average the maximum similarities across all peaks:

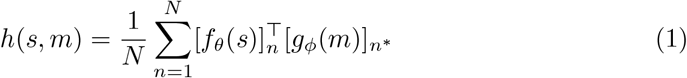

where 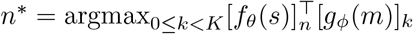 Similarly, for molecule-to-spectra similarity, each node is compared to all peaks, the maximum is taken, and we average across nodes:

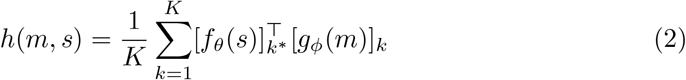

where 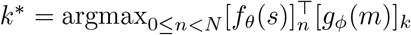.

These bidirectional fine-grained similarities replace the standard global similarity function in contrastive learning. The loss itself remains the standard symmetric contrastive loss over a batch ℬ of spectra–molecule pairs:

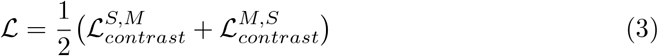

with

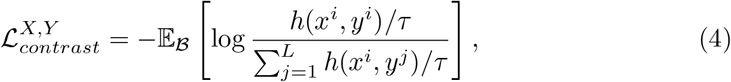

where ℬ= (*x*^1^, *y*^1^), (*x*^2^, *y*^2^), …, (*x*^*L*^, *y*^*L*^) is a batch of *L* positive pairs and *τ* is a temperature parameter.

### 4.4 Pairwise embedding similarity against fingerprint Tanimoto similarity

Pairs of spectra are sampled from MassSpecGym to ensure coverage across the full range of Tanimoto similarities. We evaluate up to 20 million randomly selected spectrum pairs and retain at most 2,000 pairs per similarity bin, using 40 equally spaced bins spanning the interval [0,1]. This procedure yielded a final set of over 50,000 spectrum pairs. Tanimoto similarity between molecules is computed by first generating Morgan fingerprints (radius = 2, nBits = 4096) and then applying the Tanimoto similarity function implemented in RDKit [34]. The FLARE spectra and molecule embeddings are computed by averaging the peak and node embeddings, respectively. Five independent runs are conducted, and the average Pearson and Spearman correlations and their standard deviations are reported (Fig. 2f).

To run MS2DeepScore, we use the publicly available checkpoint trained on more than 500,000 spectra [35]. Similarly for DreaMS, we use the publicly available model in a zero-shot manner. Embedding similarities are calculated as the cosine similarity between the respective embeddings. Modified cosine similarity is computed using the ModifiedCosine function (tolerance=0.2) from the matchms [36] library.

### 4.5 UMAP visualization of spectral embeddings

For the purpose of visualization, we include only spectra whose corresponding molecules belong to the ten most frequent classes. Class assignments were obtained from the superclass annotations provided by NPClassifier [37]. To ensure that the embedding space reflects the distribution of the original data, we sample spectra in proportion to their occurrence in the full dataset, restricting the selection to at most one spectrum per molecule. This procedure yielded a set of 13,133 spectra. Spectra embeddings are obtained by averaging peak embeddings and subsequently projected using the UMAP python package with n neighbors=45, metric=‘cosine’, min dist=0.8.

### 4.6 Class prediction using K-nearest neighbors

FLARE spectral embeddings organize tandem mass spectra into a chemically meaningful latent space in which spectra cluster by the corresponding chemical superclass. In typical untargeted metabolomics experiments, where full structural annotation is often unavailable, these embeddings enable prediction of superclass-level annotations, which are useful for many downstream analyses.

We evaluated this approach by predicting superclasses for 23 low-confidence yet statistically significant features from our case study. MassSpecGym spectra were used as reference data, restricted to the 20 most frequent chemical superclasses. Superclass prediction was performed using a k-nearest neighbors classifier (KNeighborsClassifier, scikit-learn[38]) with 20 neighbors (n neighbors = 20) and cosine distance, operating in the FLARE embedding space.

### 4.7 Unconstrained candidate retrieval via cross-modal embedding

FLARE’s cross-modal alignment enables molecular retrieval without requiring prior constraints on molecular formula or precursor mass. Because alignment is learned at a fine-grained, substructure level, spectral similarity reflects shared substructural compatibility rather than exact molecular matches. Consequently, even when no high-confidence candidates are present, embedding-based retrieval prioritizes molecules sharing chemically meaningful substructures.

We evaluate this capability by retrieving candidate molecules for the 23 features that could not be confidently annotated. All molecules from the same database used in the initial retrieval step (approximately 2 million molecules) were embedded in advance, with molecular node embeddings aggregated by averaging. For each query spectrum, we use Faiss[39] to identify the top 20 candidates by cosine similarity in the embedding space, followed by secondary re-ranking using the FLARE fine-grained similarity score. The top five candidates were retained for visualization and substructure annotation.

### 4.8 PDX sample extraction and data collection

#### 4.8.1 Sample Preparation and Metabolite Extraction of Tumor Samples

Untargeted metabolomics was performed using previously described methods [40–44]. Approximately 50 mg of snap-frozen tissue was collected from primary and metastatic breast patient-derived xenograft (PDX) tumors. Each sample was placed in a tube containing ceramic beads, and 500 µL of ice-cold homogenization solution (80% methanol, 20% water) was added. Samples were homogenized using a bead homogenizer (speed: 5.0 m/s; duration: 30 s). Homogenates were centrifuged at 4°C at 16,000 × g for 10 min. After centrifugation, 50 µL of each supernatant was transferred to a new tube and dried using a speed vacuum concentrator. Samples were reconstituted in 95% water and 5% methanol at a volume proportional to the original tissue weight. Reconstituted samples were vortexed at 2,500 rpm for 10 min and centrifuged again at 4°C at 16,000 × g for 10 min. Supernatants were transferred to LC-MS vials. Study pools were prepared by combining 10 µL from each sample. Method blanks were generated by adding 200 µL of homogenization solution to tubes with ceramic beads and processing identically to study samples.

#### 4.8.2 Untargeted Metabolomics Analysis by LC-MS

Metabolomics data were acquired using a Vanquish UHPLC system coupled to a Q Exactive™ HF-X Hybrid Quadrupole-Orbitrap Mass Spectrometer (Thermo Fisher Scientific, San Jose, CA). Metabolites were separated on an HSS T3 C18 column (2.1 × 100 mm, 1.7 µm; Waters Corporation) maintained at 50°C. The mobile phases consisted of water (A) and methanol (B), each containing 0.1% formic acid (v/v). The UHPLC gradient was as follows: 1% B for 1 min, ramp to 99% B over 16 min, hold at 99% B for 3 min, then return to 1% B for 3 min; flow rate was 400 µL/min throughout. Untargeted data were acquired in data-dependent acquisition mode over an m/z range of 70–1050. Method blanks and study pool injections were interspersed evenly throughout the study samples.

#### 4.8.3 Metabolomics Raw Data Preprocessing

Raw data were processed using Progenesis QI (version 2.1; Waters Corporation) for peak picking, alignment, and normalization. Background signals were removed by excluding features with a fold change *<* 3 in study pool versus blank injections. Normalization was performed using the “normalize to all” feature in Progenesis QI[45].

## 5 Data Availability

The MassSpecGym benchmark dataset is curated and provided by [20]. Metabolomics data from primary and metastatic breast patient-derived xenograft tumor samples are available on the Metabolomic Workbench under the study number ST003752.

## 6 Code Availability

All resources necessary to replicate our experiments, including code for training new models, loading the pretrained models, and visualizing peak-to-node correspondence, are publicly available at https://huggingface.co/spaces/HassounLab/FLARE.

## 7 Supplementary information

Supplementary Tables 1-3 and Figure 1.

## S1 Supplementary Tables and Figures

**Table S1:** MassSpecGym statistics on the whole dataset.

|  | Train | Val | Test |
| --- | --- | --- | --- |
| Spectra count | 194,119 | 19,429 | 17,556 |
| Molecule count | 25,046 | 3,386 | 3,170 |

**Table S2:** Model hyperparameter search space and selected model hyperparameters. The model was tuned using Optuna[46] with the provided search space. The final model architecture was selected based on the smallest validation loss.

| Parameter | Search Space | Selected |
| --- | --- | --- |
| Formula Linear Projection | (64,128), (512,256), (256,512), (128), (256), (128,128), (512,512), (64,64,64) | (512, 256) |
| Spectra encoder n-attention-heads | 2, 4, 8 | 4 |
| Spectra encoder n-layers | 1, 2, 3, 4, 5 | 2 |
| Molecule encoder hidden dimension | (64,128), (128,256), (256,512), (64,128,128), (128,128), (64,64,64) | (128, 256) |
| Final peak/node embedding dimension | 64, 256, 512, 1024 | 512 |
| Dropout | 0.1 - 0.5 | 0.1571042734775 |
| Contrastive temperature $\tau$ | 0.01 - 0.1 | 0.0227 |
| Learning rate | 1e-6 - 1e-3 | 2.8813e-05 |
| Batch size | 32, 64, 128, 256 | 64 |
| Weight decay | 1e-6 - 1e-2 | 1.8376e-05 |

**Table S3:**
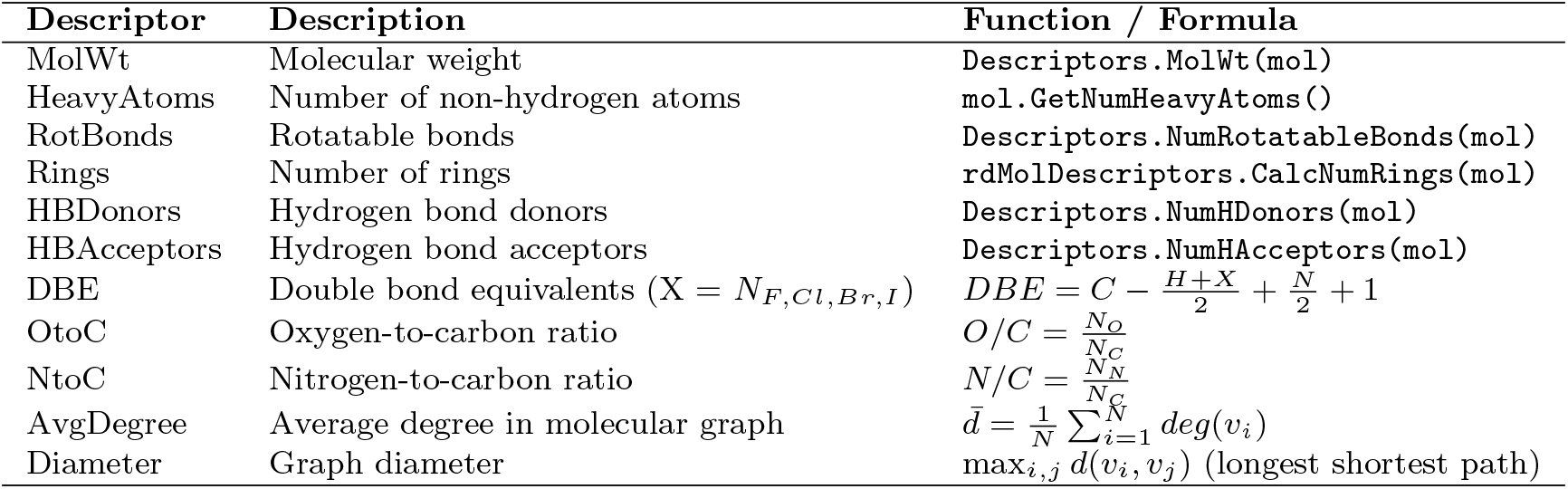
Summary of computed molecular descriptors. RDKit-based descriptors are computed using functions from rdkit.Chem.Descriptors and rdkit.Chem.rdMolDescriptors. Custom descriptors are defined by explicit formulas.

| Descriptor | Description | Function / Formula |
| --- | --- | --- |
| MolWt | Molecular weight | `Descriptors.MolWt(mol)` |
| HeavyAtoms | Number of non-hydrogen atoms | `mol.GetNumHeavyAtoms()` |
| RotBonds | Rotatable bonds | `Descriptors.NumRotatableBonds(mol)` |
| Rings | Number of rings | `rdMolDescriptors.CalcNumRings(mol)` |
| HBDonors | Hydrogen bond donors | `Descriptors.NumHDonors(mol)` |
| HBAcceptors | Hydrogen bond acceptors | `Descriptors.NumHAcceptors(mol)` |
| DBE | Double bond equivalents ($X = N_{F,Cl,Br,I}$) | $DBE = C - \frac{H+X}{2} + \frac{N}{2} + 1$ |
| OtoC | Oxygen-to-carbon ratio | $O/C = \frac{N_O}{N_C}$ |
| NtoC | Nitrogen-to-carbon ratio | $N/C = \frac{N_N}{N_C}$ |
| AvgDegree | Average degree in molecular graph | $\bar{d} = \frac{1}{N} \sum_{i=1}^N deg(v_i)$ |
| Diameter | Graph diameter | $\max_{i,j} d(v_i, v_j)$ (longest shortest path) |

**Fig. S1:**
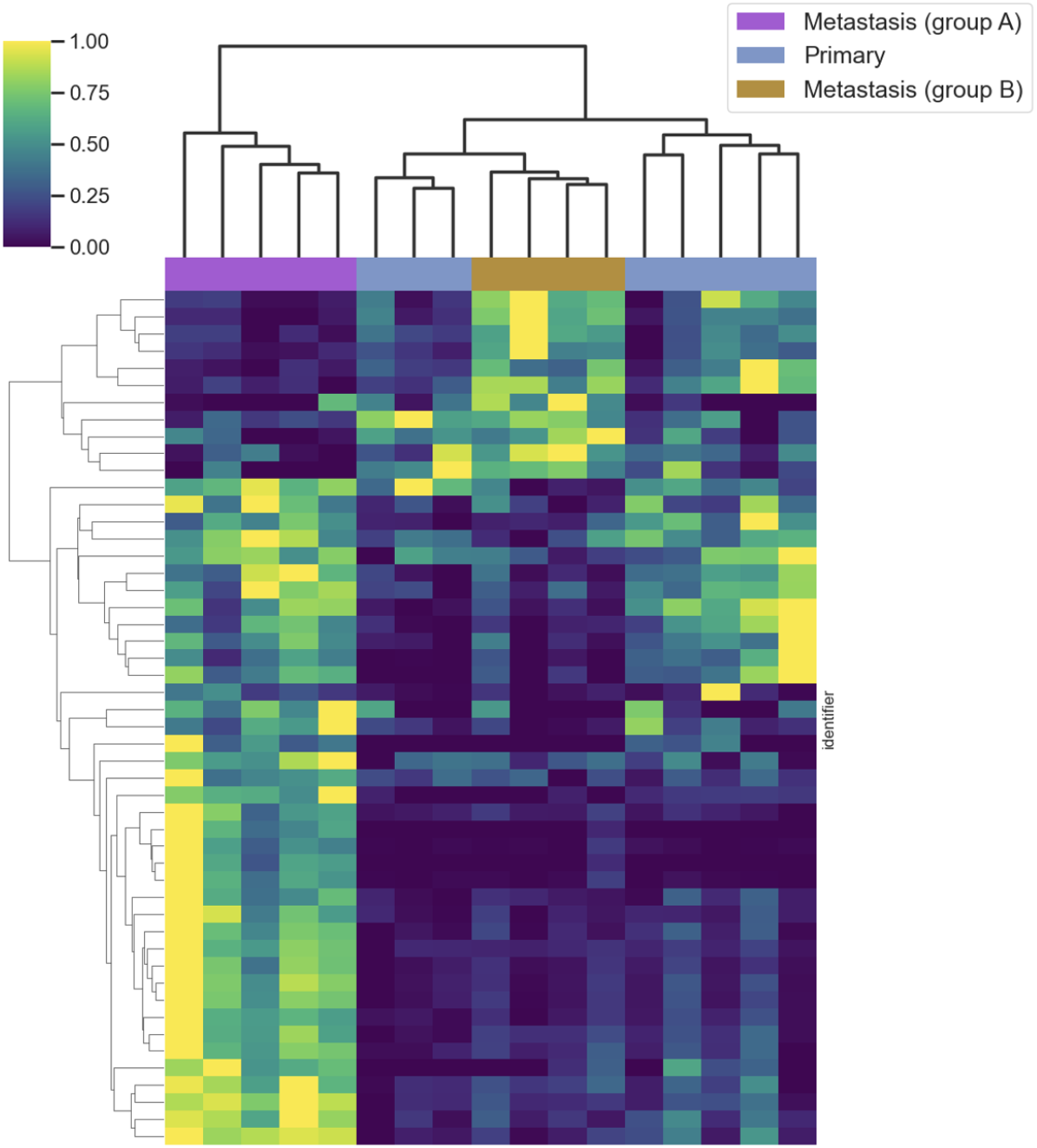
Clustered heatmap of all samples with features selected based on the 50 most significant features for differentiating samples of metastasis groups A and B.

## References

[1] Pang, H., Hu, Z.: Metabolomics in drug research and development: the recent advances in technologies and applications. Acta Pharmaceutica Sinica B 13(8), 3238–3251 (2023)

[2] Schmidt, D.R., Patel, R., Kirsch, D.G., Lewis, C.A., Vander Heiden, M.G., Locasale, J.W.: Metabolomics in cancer research and emerging applications in clinical oncology. CA: a cancer journal for clinicians 71(4), 333–358 (2021)

[3] Bittremieux, W., Wang, M., Dorrestein, P.C.: The critical role that spectral libraries play in capturing the metabolomics community knowledge. Metabolomics 18(12), 94 (2022)

[4] Zhu, H., Liu, L., Hassoun, S.: Using graph neural networks for mass spectrometry prediction. arXiv preprint 2010.04661 (2020)

[5] Li, X., Zhou Chen, Y., Kalia, A., Zhu, H., Liu, L.-p., Hassoun, S.: An ensemble spectral prediction (esp) model for metabolite annotation. Bioinformatics 40(8), 490 (2024)

[6] Goldman, S., Wohlwend, J., Stražar, M., Haroush, G., Xavier, R.J., Coley, C.W.: Annotating metabolite mass spectra with domain-inspired chemical formula transformers. Nature Machine Intelligence 5(9), 965–979 (2023)

[7] Wang, R., Manjrekar, M., Mahjour, B., Avila-Pacheco, J., Provenzano, J., Reynolds, E., Lederbauer, M., Mashin, E., Goldman, S., Wang, M., et al.: Neural spectral prediction for structure elucidation with tandem mass spectrometry. BioRxiv (2025)

[8] Nowatzky, Y., Russo, F.F., Lisec, J., Kister, A., Reinert, K., Muth, T., Benner, P.: Fiora: Local neighborhood-based prediction of compound mass spectra from single fragmentation events. Nature Communications 16(1), 2298 (2025)

[9] Dührkop, K., Fleischauer, M., Ludwig, M., Aksenov, A.A., Melnik, A.V., Meusel, M., Dorrestein, P.C., Rousu, J., Böcker, S.: Sirius 4: a rapid tool for turning tandem mass spectra into metabolite structure information. Nature methods 16(4), 299–302 (2019)

[10] Wang, F., Liigand, J., Tian, S., Arndt, D., Greiner, R., Wishart, D.S.: Cfm-id 4.0: more accurate esi-ms/ms spectral prediction and compound identification. Analytical chemistry 93(34), 11692–11700 (2021)

[11] Chen, L., Xia, B., Wang, Y., Huang, X., Gu, Y., Wu, W., Zhou, Y.: Cmssp: A contrastive mass spectra-structure pretraining model for metabolite identification. Analytical Chemistry 96(42), 16871–16881 (2024)

[12] Kalia, A., Zhou Chen, Y., Krishnan, D., Hassoun, S.: Jestr: J oint e mbedding s pace t echnique for r anking candidate molecules for the annotation of untargeted metabolomics data. Bioinformatics, 354 (2025)

[13] Xie, T., Zhang, H., Yang, Q., Sun, J., Wang, Y., Long, J., Zhang, Z., Lu, H.: Csums2: A contrastive learning framework for cross-modal compound identification from ms/ms spectra to molecular structures. Analytical Chemistry (2025)

[14] Zhou Chen, Y., Hassoun, S.: Learning from all views: A multiview contrastive framework for metabolite annotation. bioRxiv, 2025–11 (2025)

[15] Radford, A., Kim, J.W., Hallacy, C., Ramesh, A., Goh, G., Agarwal, S., Sastry, G., Askell, A., Mishkin, P., Clark, J., et al.: Learning transferable visual models from natural language supervision. In: International Conference on Machine Learning, pp. 8748–8763 (2021). PmLR

[16] Yao, L., Huang, R., Hou, L., Lu, G., Niu, M., Xu, H., Liang, X., Li, Z., Jiang, X., Xu, C.: Filip: Fine-grained interactive language-image pre-training. arXiv preprint 2111.07783 (2021)

[17] Jia, C., Yang, Y., Xia, Y., Chen, Y.-T., Parekh, Z., Pham, H., Le, Q., Sung, Y.-H., Li, Z., Duerig, T.: Scaling up visual and vision-language representation learning with noisy text supervision. In: International Conference on Machine Learning, pp. 4904–4916 (2021). PMLR

[18] Tarvainen, A., Valpola, H.: Mean teachers are better role models: Weight-averaged consistency targets improve semi-supervised deep learning results. Advances in neural information processing systems 30 (2017)

[19] Tian, Y., Krishnan, D., Isola, P.: Contrastive multiview coding. In: European Conference on Computer Vision, pp. 776–794 (2020). Springer

[20] Bushuiev, R., Bushuiev, A., Jonge, N., Young, A., Kretschmer, F., Samusevich, R., Heirman, J., Wang, F., Zhang, L., Dührkop, K., et al.: Massspecgym: A benchmark for the discovery and identification of molecules. Advances in Neural Information Processing Systems 37, 110010–110027 (2024)

[21] Scarselli, F., Gori, M., Tsoi, A.C., Hagenbuchner, M., Monfardini, G.: The graph neural network model. IEEE transactions on neural networks 20(1), 61–80 (2008)

[22] Huber, F., Burg, S., Hooft, J.J., Ridder, L.: Ms2deepscore: a novel deep learning similarity measure to compare tandem mass spectra. Journal of cheminformatics 13(1), 84 (2021)

[23] Jonge, N., Joas, D., Truong, L.-J., Hooft, J.J., Huber, F.: Reliable cross-ion mode chemical similarity prediction between ms2 spectra. biorxiv, 2024–03 (2024)

[24] Bushuiev, R., Bushuiev, A., Samusevich, R., Brungs, C., Sivic, J., Pluskal, T.: Self-supervised learning of molecular representations from millions of tandem mass spectra using dreams. Nature Biotechnology, 1–11 (2025)

[25] Bittremieux, W., Schmid, R., Huber, F., Hooft, J.J., Wang, M., Dorrestein, P.C.: Comparison of cosine, modified cosine, and neutral loss based spectrum alignment for discovery of structurally related molecules. Journal of the American Society for Mass Spectrometry 33(9), 1733–1744 (2022)

[26] Ridder, L., Hooft, J.J., Verhoeven, S.: Automatic compound annotation from mass spectrometry data using magma. Mass Spectrometry 3(Special Issue 2), 0033–0033 (2014)

[27] Wishart, D.S., Guo, A., Oler, E., Wang, F., Anjum, A., Peters, H., Dizon, R., Sayeeda, Z., Tian, S., Lee, B.L., et al.: Hmdb 5.0: the human metabolome database for 2022. Nucleic acids research 50(D1), 622–631 (2022)

[28] Sorokina, M., Merseburger, P., Rajan, K., Yirik, M.A., Steinbeck, C.: Coconut online: collection of open natural products database. Journal of Cheminformatics 13(1), 2 (2021)

[29] Böcker Group & SIRIUS Development Team: bioDB (Curated Structure Database for SIRIUS). https://drive.google.com/file/d/1kitlMkqEExMkc9EWhykERwoibBo89/view. Accessed: 2025-10-08 (2025). https://drive.google.com/file/d/1kitlMkqEExMkc9EWhykERwoibBo89/view

[30] National Institute of Standards and Technology (NIST): Tandem Mass Spectral Library. https://www.nist.gov/programs-projects/tandem-mass-spectral-library. Accessed: 2025-12-11 (2023). https://www.nist.gov/programs-projects/tandem-mass-spectral-library

[31] Global Natural Products Social Molecular Networking (GNPS) Community: MULTIPLEX-SYNTHESIS-LIBRARY-FILTERED. https://external.gnps2.org/gnpslibrary. Accessed: 2025-10-04 (2025). https://external.gnps2.org/gnpslibrary

[32] Kretschmer, F., Seipp, J., Ludwig, M., Klau, G.W., Böcker, S.: Coverage bias in small molecule machine learning. Nature communications 16(1), 554 (2025)

[33] Kipf, T.: Semi-supervised classification with graph convolutional networks. arXiv preprint 1609.02907 (2016)

[34] Landrum, G., et al.: RDKit: Open-source cheminformatics. https://www.rdkit.org. Accessed: 2025-12-11 (2006)

[35] Jonge, N.: Dual mode MS2DeepScore model (MS2DeepScore ¿2.6). https://doi.org/10.5281/zenodo.17826815. Zenodo model checkpoint, Accessed 2025-12-11 (2025). 10.5281/zenodo.17826815

[36] Huber, F., Verhoeven, S., Meijer, C., Spreeuw, H., Castilla, E.M.V., Geng, C., Hooft, J.J., Rogers, S., Belloum, A., Diblen, F., et al.: Matchms-processing and similarity evaluation of mass spectrometry data. BioRxiv, 2020–08 (2020)

[37] Kim, H.W., Wang, M., Leber, C.A., Nothias, L.-F., Reher, R., Kang, K.B., Van Der Hooft, J.J., Dorrestein, P.C., Gerwick, W.H., Cottrell, G.W.: Npclassifier: a deep neural network-based structural classification tool for natural products. Journal of natural products 84(11), 2795–2807 (2021)

[38] Pedregosa, F., Varoquaux, G., Gramfort, A., Michel, V., Thirion, B., Grisel, O., Blondel, M., Prettenhofer, P., Weiss, R., Dubourg, V., et al.: Scikit-learn: Machine learning in python. the Journal of machine Learning research 12, 2825–2830 (2011)

[39] Douze, M., Guzhva, A., Deng, C., Johnson, J., Szilvasy, G., Mazaré, P.-E., Lomeli, M., Hosseini, L., Jégou, H.: The faiss library. IEEE Transactions on Big Data (2025)

[40] Falcone, I.G., Rushing, B.R.: Untargeted metabolomics reveals acylcarnitines as major metabolic targets of resveratrol in breast cancer cells. Metabolites 15(4), 250 (2025)

[41] Rushing, B.R., Molina, S., Sumner, S.: Metabolomics analysis reveals altered metabolic pathways and response to doxorubicin in drug-resistant triple-negative breast cancer cells. Metabolites 13(7), 865 (2023)

[42] Rushing, B.R., Wiggs, A., Molina, S., Schroder, M., Sumner, S.: Metabolomics analysis reveals novel targets of chemosensitizing polyphenols and omega-3 polyunsaturated fatty acids in triple negative breast cancer cells. International Journal of Molecular Sciences 24(5), 4406 (2023)

[43] Rushing, B.R., Fogle, H.M., Sharma, J., You, M., McCormac, J.P., Molina, S., Sumner, S., Krupenko, N.I., Krupenko, S.A.: Exploratory metabolomics underscores the folate enzyme aldh1l1 as a regulator of glycine and methylation reactions. Molecules 27(23), 8394 (2022)

[44] Sharma, J., Rushing, B.R., Hall, M.S., Helke, K.L., McRitchie, S.L., Krupenko, N.I., Sumner, S.J., Krupenko, S.A.: Sex-specific metabolic effects of dietary folate withdrawal in wild-type and aldh1l1 knockout mice. Metabolites 12(5), 454 (2022)

[45] Välikangas, T., Suomi, T., Elo, L.L.: A systematic evaluation of normalization methods in quantitative label-free proteomics. Briefings in bioinformatics 19(1), 1–11 (2018)

[46] Akiba, T., Sano, S., Yanase, T., Ohta, T., Koyama, M.: Optuna: A next-generation hyperparameter optimization framework. In: Proceedings of the 25th ACM SIGKDD International Conference on Knowledge Discovery & Data Mining, pp. 2623–2631 (2019)

